# Continuous Light-Induced Circadian Rhythm Disruption Impairs Intestinal Barrier Integrity in Male C57BL/6 Mice Through Gut Microbiota Dysbiosis and the Apoptosis-Inflammation-Oxidative Stress Cascade

**DOI:** 10.1101/2025.11.11.687914

**Authors:** Tai-Wei Zhang, Jia-Chun Song, Ning-Bo Hao, Ming-Yue Qu, Bao-Shi Guo, Chang-Zheng Li

## Abstract

**Background:** Circadian rhythm disruption (CRD) is a risk factor for irritable bowel syndrome (IBS), but the mechanism linking CRD to intestinal barrier dysfunction remains unclear. This preclinical study aimed to clarify whether CRD impairs intestinal barrier integrity via gut microbiota dysbiosis and the “apoptosis-inflammation-oxidative stress” cascade.

**Methods:** Twenty-four male C57BL/6 mice were randomized into control (12h light/dark, n=12) and CRD (21-day continuous light, n=12) groups. Circadian disruption was verified via locomotor activity, serum melatonin/serotonin, and clock gene expression. Intestinal barrier function, microbiota, apoptosis, inflammation, and oxidative stress were assessed using FITC-dextran permeability, 16S rRNA sequencing, Western blotting (WB), TUNEL, and ELISA.

**Results:** CRD increased intestinal permeability (+114.7%, p<0.001), shortened villi (−25.6%, p=0.018), downregulated tight junction proteins (ZO-1, Occludin, p<0.05), and altered microbiota (family-level: decreased Prevotellaceae, increased Bacteroidaceae, p<0.05). It also activated the apoptosis-inflammation-oxidative stress cascade (Caspase-3/β-actin: +1.4-fold, IL-1β: +44.1%, MDA: +50%, CAT: −90%, all p<0.05).

**Conclusions:** CRD impairs intestinal barrier integrity via gut microbiota dysbiosis and the apoptosis-inflammation-oxidative stress cascade. These preclinical findings identify gut microbiota and the apoptosis-inflammation-oxidative stress cascade as potential targets for further investigating IBS associated with CRD.

## 1. Introduction

The circadian rhythm regulates various physiological functions, including sleep-wake cycles, metabolism, and gastrointestinal activities. These endogenous rhythms are generated through a transcription-translation feedback loop formed by core clock genes such as clock (Clock), BMAL1, PER, and CRY, which synchronize with environmental signals (particularly illumination)^[1, 2]^. This sophisticated regulatory system becomes compromised when disrupted by shift work, prolonged light exposure, or sleep pattern disturbances, which has been shown to increase the risk of cardiovascular diseases, metabolic syndrome, and neuropsychiatric disorders^[3–6]^.

Functional gastrointestinal disorders (particularly irritable bowel syndrome [IBS]) are closely associated with circadian rhythm disruptions. Epidemiological data indicate that IBS patients frequently experience sleep disturbances, irregular schedules, or night shift work patterns^[7]^. Recent experimental studies have confirmed that circadian rhythm disorders can induce IBS-like clinical manifestations and alter metabolic markers in mouse feces^[8]^. Furthermore, clinical research has revealed that individuals with gut-brain interaction dysfunction exhibit melatonin metabolism abnormalities and decreased circadian hormone levels^[9]^,because the circadian rhythm is crucial for regulating gastrointestinal physiology, any disorder may lead to gastrointestinal (GI) dysfunction. The disorders of gut-brain interaction (DGBI), a functional gastrointestinal disorder. highlighting the potential mechanistic link between circadian rhythm disorders and intestinal dysfunction.

The intestinal epithelial barrier serves as the body’s defense mechanism, acting as a dual defense system that blocks gut antigens and pathogens while regulating nutrient absorption. Its integrity depends on tight junction proteins (e.g., ZO-1, Occludin, Claudins) and is closely associated with gut microbiota that modulate immunity and metabolism^[10]^. Although circadian rhythm disruption has been shown to alter microbial composition^[11]^, it remains unclear whether these factors increase intestinal permeability through a cascade of events involving epithelial cell apoptosis, inflammation, and oxidative stress. To address this gap, we established a circadian rhythm disruption model in mice using continuous light exposure^[12]^. The study systematically evaluated behavioral rhythms, neuroendocrine factors (melatonin and serotonin), whole-body circadian clock gene expression, gut microbiota composition, and barrier integrity biomarkers, The possible mechanisms related to apoptosis, inflammation and oxidative stress were also studied. This research aims to elucidate how chronic circadian rhythm disruption impairs intestinal barrier function, providing insights into the pathogenesis of gastrointestinal disorders such as irritable bowel syndrome (IBS).

## 2. Materials and methods

### 2.1 Animal models and experimental design

Specific pathogen-free (SPF) male C57BL/6JNifdc mice (total number: 24) were selected (License No.: SCXK[SU]2021-0005, issued by Suzhou Xishan Biotechnology Co., LTD), and they were raised for one week before the experiment. Temperature 22±1℃, humidity 50±10%, The light-dark cycle was set at 12 hours of light followed by 12 hours of darkness (light intensity>350 lux, dark period <1 lux, measured using a digital light meter, Model: XY-1309, Manufacturer: Qingdao Xinye Environmental Protection Technology Co., LTD), with mice freely consuming standard feed and sterile water. After the adaptation period, they were randomly divided into two groups (n=12) using a random number table: (1) Normal circadian rhythm control group (Con group): maintained a standard 12-hour light-dark cycle (light period 08:00–20:00, dark period 20:00–08:00); (2) Circadian rhythm disruption group (CRD group): subjected to continuous light intervention (24 hours/day) for 21 days. All mice were placed in the Comprehensive Laboratory Animal Monitoring System (CLAMS Columbus Instruments, OH, USA) to record feeding, autonomous activity, and wheel exercise parameters in real time.

### 2.2 Sample collection and processing

Each group was randomly sampled with three mice at time points 08:00,14:00,20:00, and at 02:00 of the next day to ensure a minimum sample size with statistically significant data^[13]^. Euthanasia was performed via cervical dislocation. To precisely evaluate the impact of circadian rhythm disruption on hormone levels, we collected samples at 12 time points during dynamic analysis of serum melatonin and serotonin (5-HT). Blood, hypothalamic tissue, and ileal tissues were immediately collected post-destruction (using ice preservation). Serum was obtained through centrifugation, while tissue samples were either rapidly frozen in liquid nitrogen and stored at −80℃ or fixed in 4% polyformaldehyde solution (Biosharp) for later use.

### 2.3 Western blotting (WB)

The ice-cold grinding of samples was conducted in RIPA lysis buffer (Beyotime) containing protease inhibitors (AbMole BioScience) at a 1:100 dilution for 30 minutes. After lysis, the samples were centrifuged at 4°C at 14,000×g for 15 minutes, followed by collection of the supernatant. Protein concentration was measured using the BCA Protein Assay Kit (Vazyme) method, then RIPA lysis buffer/SDS loading buffer (Biosharp) was added for normalization and heating to denature proteins. A 10% SDS-PAGE Gel Kit (Vazyme) gel electrophoresis was performed at 120 V for 90 minutes. Trans-Blot Turbo transfer system (Bio-Rad) was used to transfer proteins onto 0.2 μm Trans-Blot Turbo Mini PVDF membrane (Bio-Rad). The membrane was blocked with 5% skim milk in TBST at room temperature for 1.5 hours, followed by three TBST washes (15 minutes per wash). Primary antibodies (Primary antibodies are listed in Supplementary Table 1, with detailed information on sources and catalog numbers.) were incubated overnight at 4°C. The membrane was washed three times with TBST (15 minutes per wash), followed by a 1-hour room temperature incubation with HRP-labeled secondary antibody (Proteintech) at 1:5000 dilution. Finally, the membrane was washed three times with TBST (15 minutes per wash) before ECL chemiluminescence detection.

### 2.4 Enzyme-Linked Immunosorbent Assay (Elisa)

The concentrations of IL-6 (RXW203049M), TNF-α (RXM202412M), and IL-1β (RXW203076M) in tissues were determined using ELISA kits sourced from RUIXIN BIOTECH. Serum melatonin levels were evaluated with an ELISA kit acquired from Sangon Biotech (D721190). Serum 5-HT concentrations were measured using an ELISA kit from Elabscience (E-EL-0033), in accordance with the manufacturer’s protocols. Serum samples were diluted 5-fold prior to analysis. Tissue homogenates underwent centrifugation to harvest the supernatant, which was then diluted 10-20 times with dilution buffer. The absorbance (OD value) for each well was recorded at 450 nm wavelength using a microplate reader.

### 2.5 Detection of oxidative stress indicators

Total Superoxide Dismutase (SOD) activity (Bioengineering product code A007-1-1): Assessed using the NBT reduction method. Reduced Glutathione (GSH-PX) content (Bioengineering product code A005-1-2): Measured through the DTNB colorimetric assay. Malondialdehyde (MDA) content (Bioengineering product code A003-1-2): Evaluates lipid peroxidation levels using the thiobarbituric acid (TBA) method. Catalase (CAT) activity (Bioengineering product code A001-3-2): Analyzed via the ammonium molybdate termination method.

### 2.6 Behavioral experiment (open field experiment)

The experiment utilized a 40×40×35 cm transparent acrylic enclosure with 20 lux illumination and noise levels below 50 dB (as per Zhongshi Technology). The procedure consisted of three phases: an adaptation phase (conducted independently for 30 minutes), a testing phase (video recording for 5 minutes), and a cleaning phase (using 75% ethanol for wiping). Data collection and analysis were performed using Tracking Master V5 software.

### 2.7 FITC-dextran in vivo imaging and serum FITC-dextran detection

The FITC-dextran (FD4,60842-46-8, SIGMA-ALDRICH, average molecular weight 3000-5000) was weighed and dissolved in sterile PBS (pH 7.4) as a pale yellow powder at a final concentration of 60 mg/mL (dissolved by vortex-shaking under light protection). Gavage administration (Day 21 post-modeling): After fasting all mice for 4 hours (with free access to water), the hair was removed from their abdomens. FITC-dextran solution was administered via oral gavage at 600 mg/kg body weight. Three to four hours after administration, 10% chloral hydrate was intraperitoneally injected for anesthesia. The animals were then placed in a dark chamber small animal in vivo imaging system (IVIS Spectrum, PerkinElmer) with the following parameters: excitation/emission wavelengths set at 490 nm and 525 nm respectively, capturing fluorescence signals from the abdominal region in a supine position.

Immediately after live imaging, eyeball blood was collected and allowed to stand at room temperature (protected from light) for 30 minutes. The serum was separated by centrifugation at 13,000×g for 15 minutes at 4°C, followed by collection of the supernatant.Dilute the serum with PBS at a 1:5 ratio (linear range confirmed in preliminary experiments). Add 100μL of diluted serum to a black 96-well plate and perform the assay using a microplate reader with excitation wavelength set at 485 nm and emission wavelength at 528 nm. dilution concentration is specified in Supplementary Table 2.

### 2.8 H&E stain

The tissue was fixed with 4% PFA (Biosharp) for 24 hours. Dehydration and embedding: Treated with graded ethanol (MACKLIN, 70%-100%) for dehydration, xylene (Millipore) for transparency, and paraffin (58-60℃) for embedding. Sections were cut to 4μm thickness perpendicular to the intestinal longitudinal axis. De-waxing and hydration: Xylene (two applications, 5-10 minutes each), anhydrous ethanol (two applications, 2 minutes each), 95% and 80% ethanol (two applications each), followed by 1-minute running water rinse. Subsequent staining: 3-5 minutes in hematoxylin solution (Servicebio), 1-minute running water rinse, 1-3 seconds alcohol fixation with 1% hydrochloric acid, and 5-10 minutes counterstaining with hematoxylin (Servicebio). Dehydration and transparency: 80% ethanol quick pass, 95% ethanol (two applications, 1 minute each), anhydrous ethanol (two applications, 2 minutes each), and xylene (two applications, 5 minutes each). Mounting: Sealed with neutral resin (Biosharp) under a bubble shield.

### 2.9 Small intestinal histiocytosis detection (TUNEL staining)

Conventional paraffin embedding and sectioning were performed. After dewaxing and hydration, 20μg/mL DNase-free protease K (Beyotime ST533) was added dropwise, followed by 20-minute incubation at 37℃. Sections were washed with PBS (ZSGB-BIO) and incubated in a dark chamber at 37℃for 60 minutes using TUNEL reaction mixture. Post-incubation washing with PBS terminated the reaction, followed by mounting with DAPI-containing anti-fluorescence quenching (Beyotime) mount medium.

### 2.10 16S rRNA gene sequencing

DNA samples were analyzed using UV spectrophotometry (concentration ≥2 ng/μL, A260/A280 = 1.8–2.0) and 1% agarose gel electrophoresis to verify integrity. The 16S rRNA V3–V4 region was amplified using primers 341F (5′-CCTACGGGNGGCWGCAG-3′) and 805R (5′-GACTACHVGGGTATCTAATCC-3′) via KAPA high-fidelity amplification. The PCR protocol included: 95℃ 180s pre-denaturation; 98℃ 20s,55℃ 15s,72℃ 15s,30 cycles; 72℃ 60s final extension. Amplified products were quantified by Qubit (5.839 ng/μL) and subjected to library preparation with 3’ end “A” tailing, Adapter ligation, gel recovery of target fragments (main peak 598bp confirmed by Agilent 2100, close to theoretical 465 bp), and PCR enrichment. Libraries were diluted to 12pM for paired-end sequencing on HiSeq platform (500bp read length). During the bioinformatics analysis, Sequences were filtered to remove reads with quality scores <20, ambiguous bases, or lengths outside 435-495bp. Chimeric sequences were removed using UCHIME. Clustering of Operational Taxon Units (OTUs) was performed using the Silva v132 database with a 97% similarity threshold. Diversity indices including Shannon index, ACE index, and othersα were calculated, with βdiversity visualization achieved through Principal Component Analysis (PCA), Principal Component Analysis (PCoA), and Non-Metric Distance Scaling (NMDS). Statistical analysis employed ANOSIM similarity analysis (R=0.472, p=0.002), LEfSe at the family level, and Tukey’s test/t-test for species differences between groups. The results were presented as OTU tables, community structure diagrams, and differential species lists.

### 2.11 statistical analysis

Data are presented as mean ± standard error (SEM). Sample size (n=12 per group) was calculated using G*Power 3.1 software with an α error probability of 0.05, power (1-β) of 0.8, and an expected effect size of 0.8 based on preliminary experiments. Inter-group comparisons were performed using unpaired t-tests (data met the normal distribution assumption, verified via Shapiro-Wilk test). All statistical analyses were conducted using GraphPad Prism 9.5.0 software. Significance levels were set at *p<0.05, **p<0.01, and ***p<0.001. No samples or animals were excluded from the analysis.

## 3. Result

### 3.1 CRD induced systemic physiological rhythm disorder and metabolic phenotype

The CRD group mice showed abnormal weight gain. From day 7 of the experiment, the weight of the CRD group continued to increase (Figure 1A), Significant statistical difference on day 21 (p <0.05) (Figure 1B). The difference in weight did not stem from changes in food intake (Figure 1C), There was no statistical difference in the average daily food intake between the two groups one week before modeling (p> 0.05) (Figure 1D).

**Figure 1.**
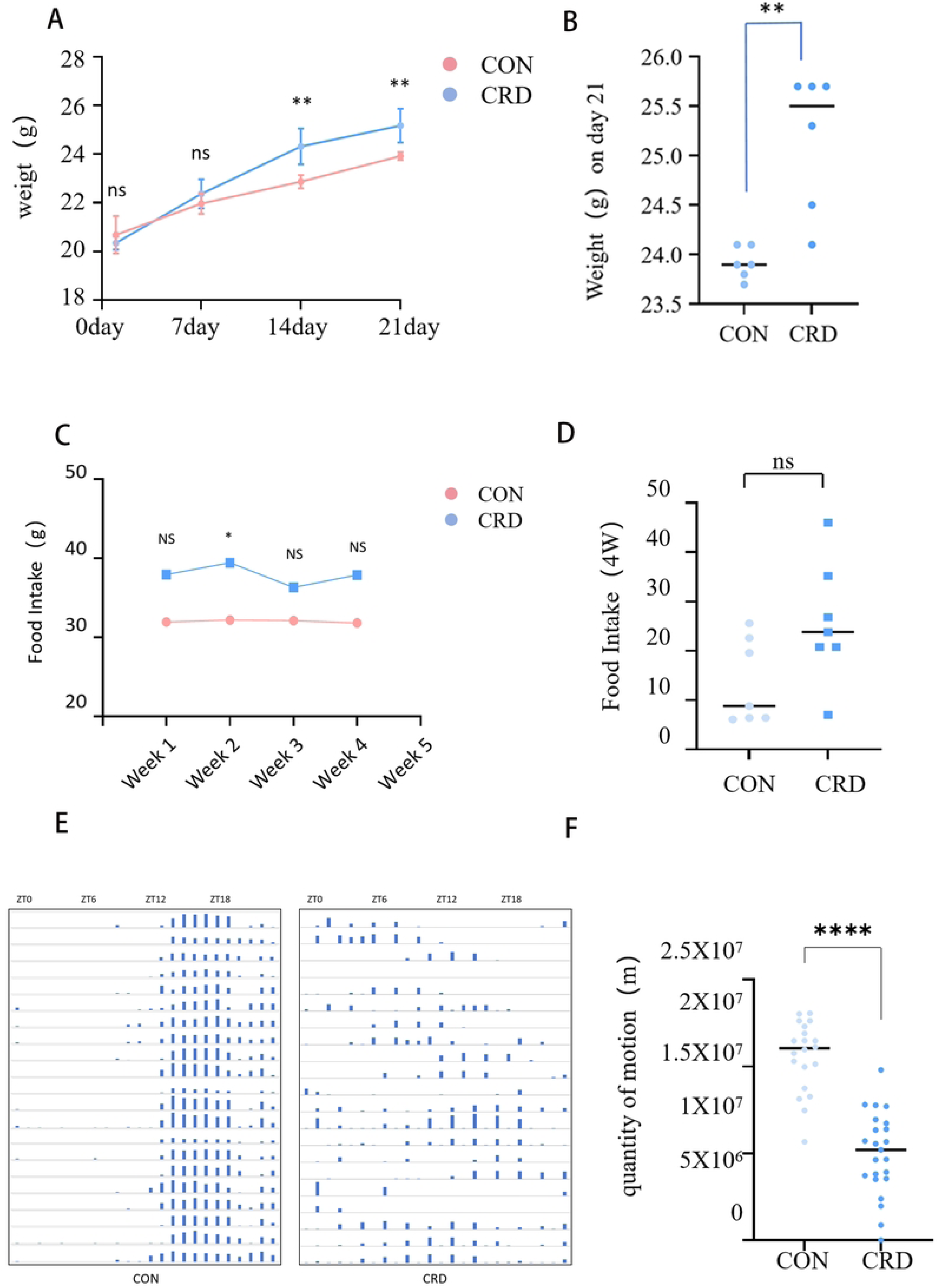
CRD influences body weight, food intake, and voluntary exercise in mice. **(A)** Body weight progression of CON and CRD groups over the 21-day experimental period. **(B)** Final body weight measured at day 21. **(C)** Average weekly food intake for both groups throughout the study. **(D)** Average daily food intake during the final w eek (days 15-21). **(E)** Running wheel activity profiles (e.g., revolutions per day) for C ON and CRD mice. **(F)** Statistical analysis of daily voluntary movement activity over the 21-day period. Data are presented as mean ± SEM(n=6/group). Statistical significance was determined by unpaired Student’s t-test (A,B,C,D,F). *p < 0.05, **p < 0.01, ** *p < 0.001 vs. CON group. control group (CON), Disruption of circadian rhythms group (CRD).

21-day running monitoring showed that CRD disrupted the loss of circadian coordination of motor behavior in mice (Figure 1E). The specific manifestations included reduced rhythmic intensity (p<0.01), loss of phase stability (p<0.001), fragmented activity patterns (p<0.01), and a 62% decrease in maximum duration. The proportion of dark-phase activity dropped to 38.5% (p<0.001), accompanied by an abnormal daytime activity peak. CRD group mice exhibited decreased total exercise distance. Exercise volume in the CRD group plummeted significantly compared to the CON group (p<0.001) (Figure 1F).

### 3.2 Behavioral and endocrine dysfunction

Open field experimentsuggested that CRD significantly inhibited spontaneous movement behavior in mice (Figure 2A), manifested as a decrease in average speed (p=0.002) (Figure 2B), Total exercise distance decreased (p <0.001) (Figure 2C), The exploratory behavior decreased significantly, including the decrease in central zone dwell time (p=0.012) (Figure 2D)and stagnate time increase(p=0.008) (Figure 2E), Behavior patterns tend to linger around.

**Figure 2.**
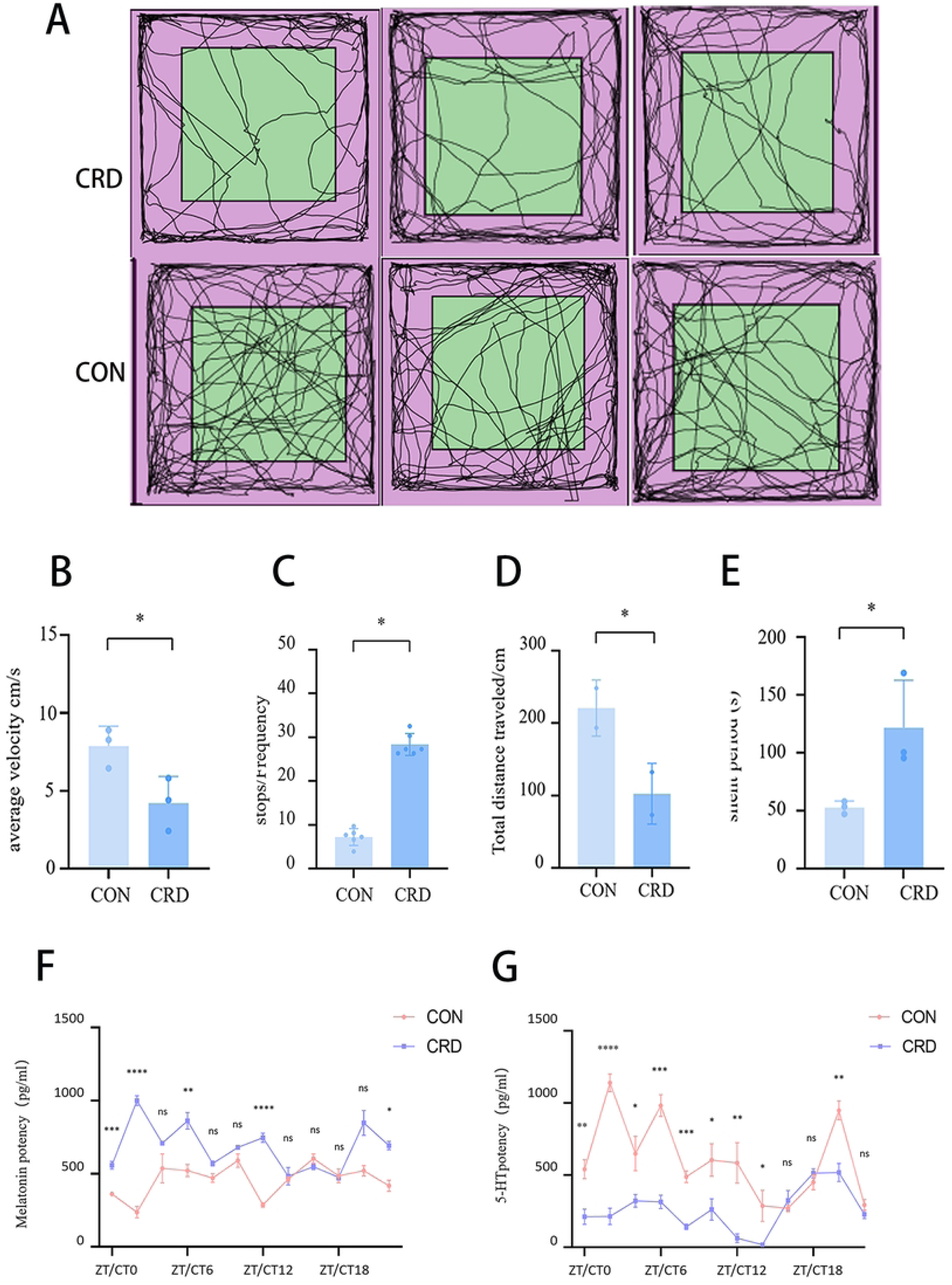
CRD suppresses spontaneous locomotor activity and disrupts circadian rhythms of melatonin and serotonin in mice. **(A)** Representative actograms from the open field test for CON and CRD groups. **(B)** Average moving speed. **(C)** Total move ment distance. **(D)** Time spent in the central zone. **(E)** Immobility time. **(F)** Circadian rhythm of serum melatonin (MT) concentrations. Blood samples were collected every 2 hours. **(G)** Circadian rhythm of serum serotonin (5-HT) concentrations. All data are presented as mean ± SEM(n=3). Statistical significance for panels B-G was determined by unpaired Student’s t-test (p-values are indicated on the graphs). *p < 0.05, **p < 0.01, ***p < 0.001 vs CON group. CON group. control group(CON), Disruption of circadian rhythms group(CRD).

In the neuroendocrine study of melatonin (MT, Figure 2F) and 5-hydroxytryptamine (5-HT, Figure 2G), The overall secretion level of CRD group was significantly lower than that of CON group (p<0.05), and the change of secretion rhythm showed phase delay and amplitude attenuation.

### 3.3 Clock gene dysfunction

In the hypothalamus(Figure 3A), CLOCK expression in CT12 was significantly reduced in the CRD group (p <0.001), suggesting impaired normal circadian accumulation (Figure 3B); CRY peaked in the CON group at ZT12, while CRD showed an abnormally high level at CT18 (p <0.001)(Figure 3C); The peak of BMAL1 in CON group ZT12 completely disappeared in CRD group (p <0.001), and showed abnormal high in CT18 (p <0.001), showing close phase reversal(Figure 3D); PER1 was significantly elevated in CRD group CT0 and CT6 (p <0.05), but significantly suppressed in ZT12 (p <0.01)(Figure 3E).

**Figure 3.**
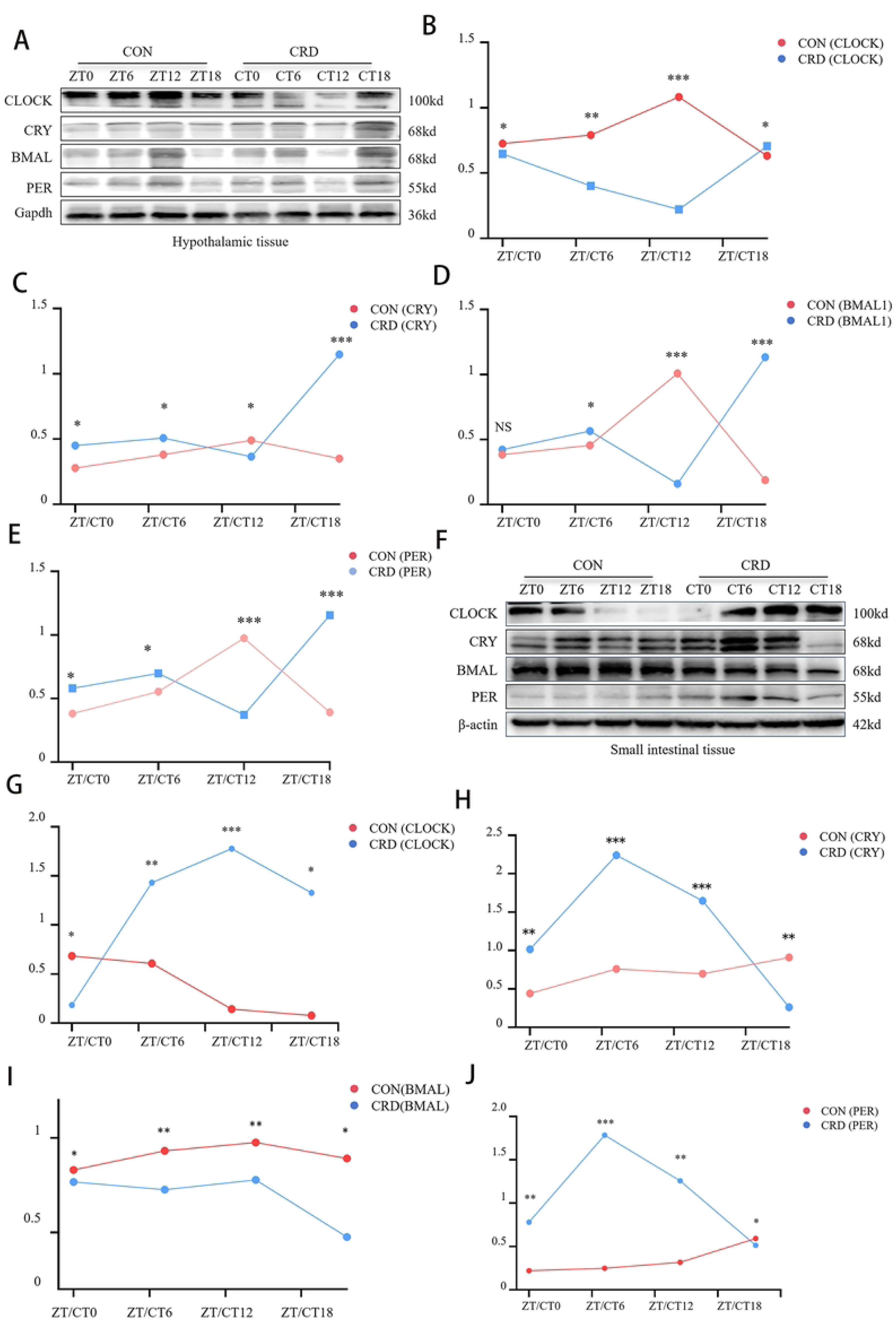
Altered expression patterns of core circadian clock proteins in the hypothalamus and small intestine of CRD mice. **(A)** Representative Western blots of CLOCK, CRY, BMAL1, and PER1 in hypothalamic tissues from control (CON, Zeitgeber Time [ZT]) and CRD (Circadian Time [CT]) mice. **(B-E)** Quantification of hypothalamic proteins: CRD altered diurnal patterns of CLOCK (p < 0.001), CRY (p < 0.001), BMAL1 (p < 0.001), and PER1 (p < 0.05) vs. CON, analyzed by unpaired Student’s t-test. **(F)** Representative Western blots of CLOCK, CRY, BMAL1, and PER1 in small intestinal tissues. **(G–J)** Quantification of small intestinal proteins: CRD induced abnormal expression of CLOCK (p < 0.05), CRY (p < 0.001), BMAL1 (p < 0.05), and PER1 (p < 0.05) vs. CON, analyzed by unpaired Student’s t-test. Data are mean ± SEM; p < 0.05, p < 0.01, *p < 0.001 vs. CON at corresponding time points (unpaired Student’s t-test).

In the small intestine, CRD also showed significant phase disruption(Figure 3F). CLOCK was significantly higher in CT12 than in CON ZT12 (p <0.05) (Figure 3G); CRY was significantly increased at CT6 (p <0.001) and decreased at CT18 (p <0.001)(Figure 3H); BMAL1 was significantly elevated at CT12 (p <0.05), and the rising trend in CON group at ZT18 turned to a decrease in CRD group(Figure 3I), Indicates that the phase of the rhythm is disrupted across tissues; PER showed a significant peak in CT6 (p <0.05), while the inherent peak of ZT18 in CON group was significantly weakened in CRD group (Figure 3J).

### 3.4 Differences in bacterial flora

At the phylum level, the relative abundance heatmap (Figure 4A) of fecal microbiota revealed similar overall composition between CON and CRD groups. Bacteroidetes, Firmicutes, and Verrucomicrobia were dominant phyla with no statistically significant differences between groups (all P> 0.05).

**Figure 4.**
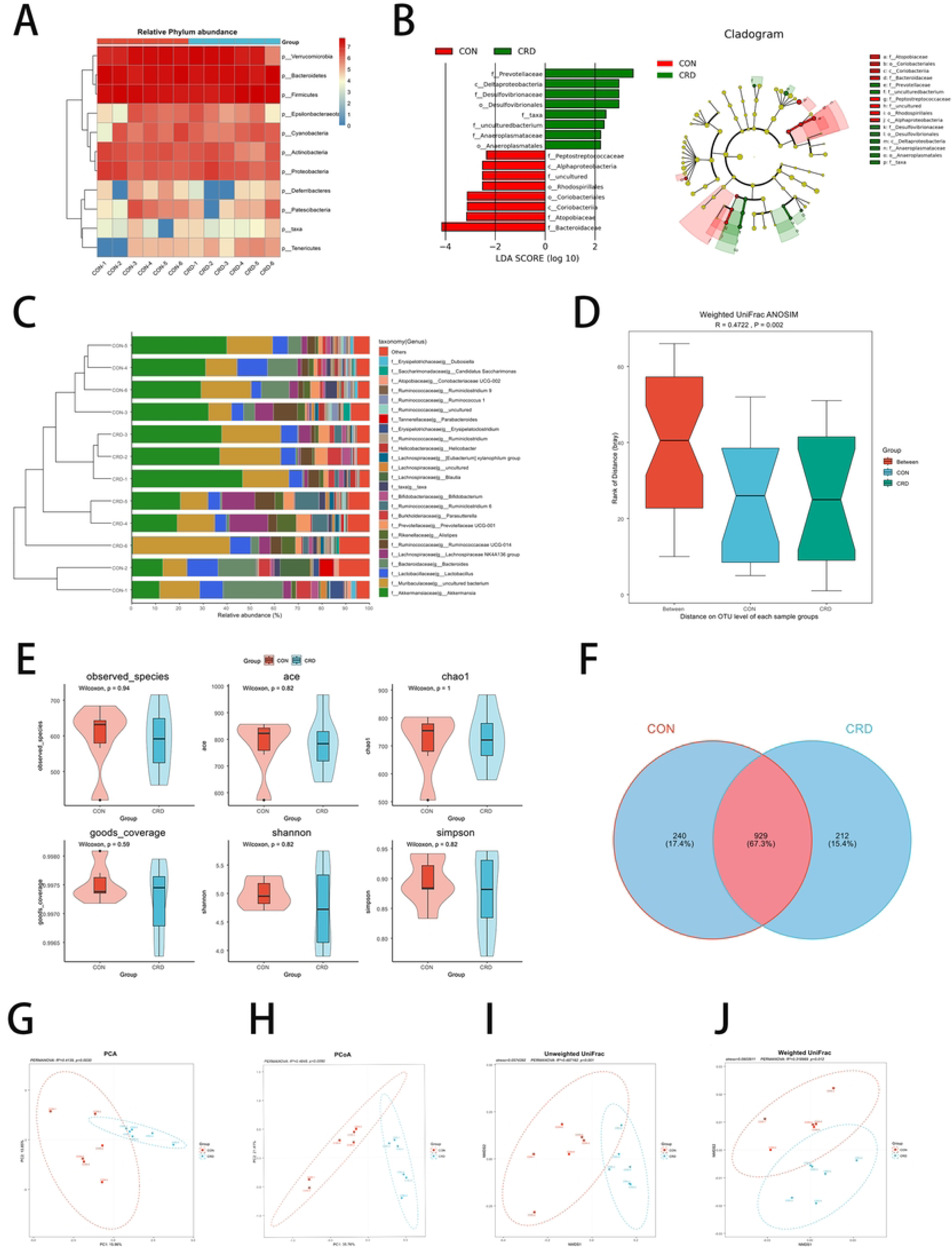
Fecal microbiota differences between the control group (CON) and circadian rhythm-disordered group (CRD). (A) Relative abundance heatmap at the phylum level; (B) Linear discriminant analysis (LEfSe) identifies differentially abundant taxa at higher taxonomic levels; (C) Species-level heatmap reveals distinct abundance patterns; (D) ANOSIM analysis (based on weighted UniFrac distances) demonstrates significant group differences (R = 0.472, P = 0.002); (E) Alpha diversity indices (Shannon index, Chao1 index, etc.) show no significant group differences (Wilcoxon test, all P> 0.05); (F) Network diagram illustrates microbial heterogeneity; (G–J) β diversity analyses: principal component analysis (PCA), principal coordinate analysis (PCoA), and non-metric multidimensional scaling (NMDS) all reveal distinct microbial profiles between CON and CRD groups.

Linear discriminant analysis (LEfSe) further identified significant taxonomic differences at a more refined classification level: CON group showed enrichment of Bacteroidaceae and Coriobacteriia (LDA score> 2, P <0.05), while CRD group exhibited specific enrichment of Prevotellaceae and δ-proteobacteria (LDA score> 2, P <0.05) (Figure 4B).

At the genus level, Akkermansia and other genera showed higher abundance in some CON samples, whereas Prevotellaceae UCG-001 and Alistipes (Bacteroidetes-branching bacteria) were more enriched in some CRD samples (Figure 4C). ANOSIM analysis for intergroup significance (Figure 4D) demonstrated significant differences with R=0.472 and p=0.002.

Comparison of diversity index: 1.Shannon index CON group showed stronger stability (standard deviation 0.23 compared with CRD group 0.64) (Figure 4E). Distribution characteristics of OTU: The Venn diagram (Figure 4F) reveals that 67.3% of OTUs are shared between the two groups, while 15.4% are unique to the CRD group, indicating higher microbial heterogeneity in the CRD group. 2. β diversity analysis: The PCA 2D plot shows a significant separation between CON and CRD samples (p=0.003), with PC1 (13.85%) and PC2 (15.96%) jointly explaining 30% of the variance (Figure 4G); PCO A results are consistent with PCA, and the two groups form independent clusters (p=0.005) (Figure 4H); NMDS analysis showed intergroup separation and intra-group clustering (Figure 4I-J).

### 3.5 CRD and intestinal barrier integrity

From a macroscopic perspective, FITC-dextran were imaged in vivo to show that: The abdominal region of CRD group showed diffuse strong fluorescence accumulation(Figure 5A), This was in sharp contrast to the weak local signals observed in the normal rhythm group (CON). Quantitative analysis confirmed that CRD group fluorescence intensity was higher than CON group(p < 0.001)(Figure 5B). The serum FITC-dextran concentration quantification corresponding to in vivo imaging was higher in CRD group than in normal rhythm group (CON) (p <0.001) (Figure 5C). H&E staining of intestinal histopathology (Figure 5D). Villon length shortened (p=0.018)(Figure 5E). Inflammatory infiltration depth increased (p=0.002() Figure 5F). crypt depth was reduced by 19.6% (p=0.018() Figure 5G). Expression of tight junction protein (TJ) at the micro level (Figure 5H). In CRD group, the expression of ZO-1, Occludin, Claudin-1 and Claudin-3 decreased (all p <0.05)(Figure 5I-L), Intestinal mechanical barrier integrity is compromised.

**Figure 5.**
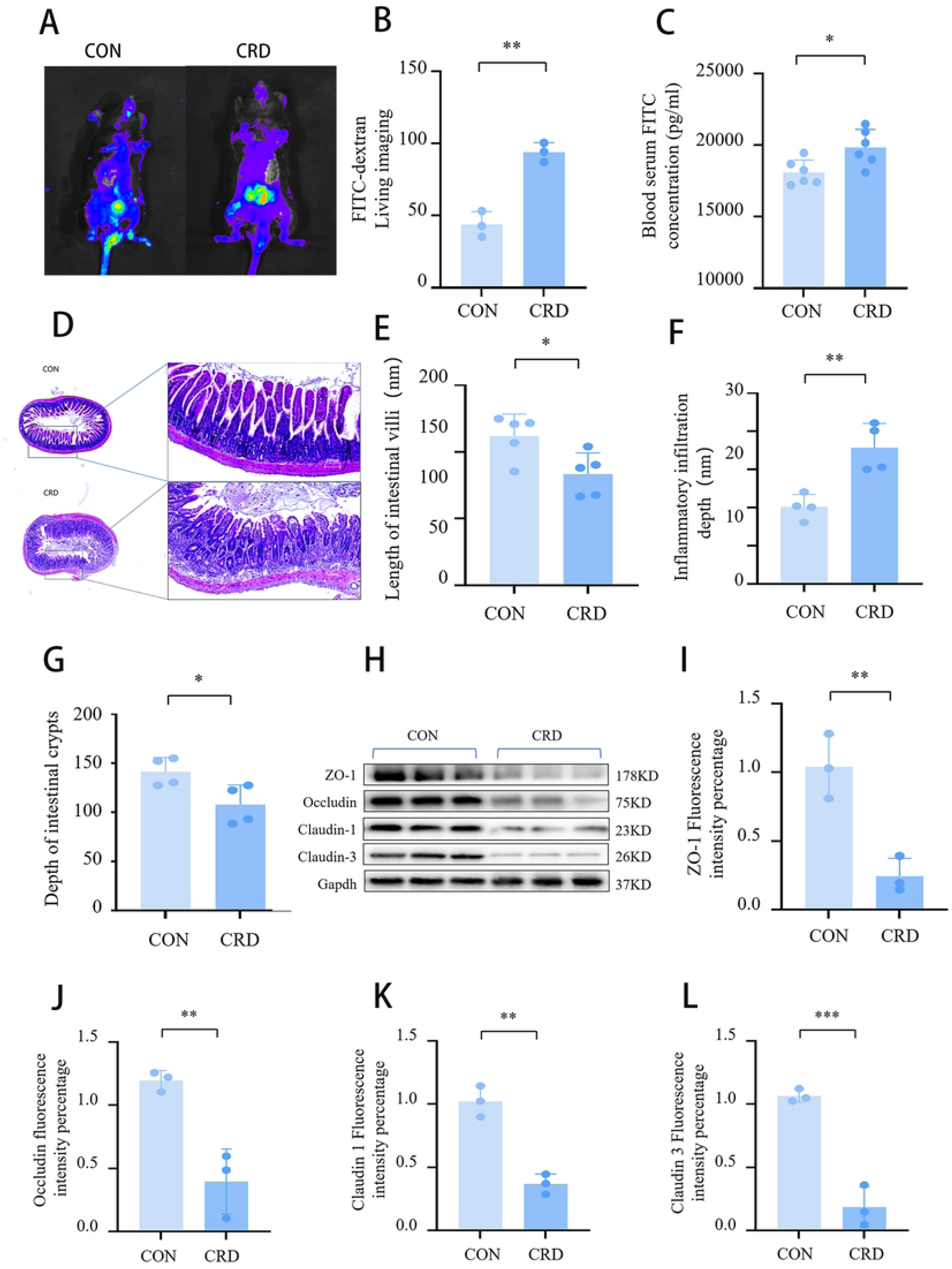
CRD disrupts intestinal barrier integrity. (A) In vivo fluorescence imaging of FITC-dextran permeability. (B) Quantification of abdominal fluorescence intensity. (C) Serum concentration of FITC-dextran. (D) Representative H&E-stained sections of small intestinal tissue(scale bar = 100 μm). CRD induced structural damage. (E) Quantification of villus length. (F) Measurement of inflammatory infiltration depth. (G) Depth of intestinal crypts. (H) Representative western blot of tight junction proteins. (I) Grey scale analysis of ZO-1, (J) Grey scale analysis of Occludin, (K) Grey scale analysis of Claudin-1, (L) Grey scale analysis of Claudin-3. All data are presented as mean ± SEM(n=3). Statistical significance was determined by unpaired Student’s t-test (p-values are indicated on the graphs). *p < 0.05, **p < 0.01, ***p < 0.001 vs CON group. CON group. control group(CON), Disruption of circadian rhythms group (CRD).

### 3.6 CRD triggers the “apoptosis-inflammation-oxidation” cascade reaction pathway of intestinal barrier damage

The expression of key apoptosis proteins (Caspase-3, Bax, Bcl-2) was detected(Figure 6A), The TUNEL fluorescence staining method was used for verification(Figure 6D). CRD group Caspase-3 significantly increased (t = 8.72, p <0.001)(Figure 6B). CRD group showed increased Bax/Bcl-2 ratio (t=6.94, p=0.002) (Figure 6C).TUNEL fluorescence staining showed that the proportion of TUNEL-positive cells in the CRD group (mean ± SEM: 18.5 ± 2.1%) was significantly higher than that in the CON group (3.2 ± 0.5%, t=4.87, p=0.0013) (Figure 6E).

**Figure 6.**
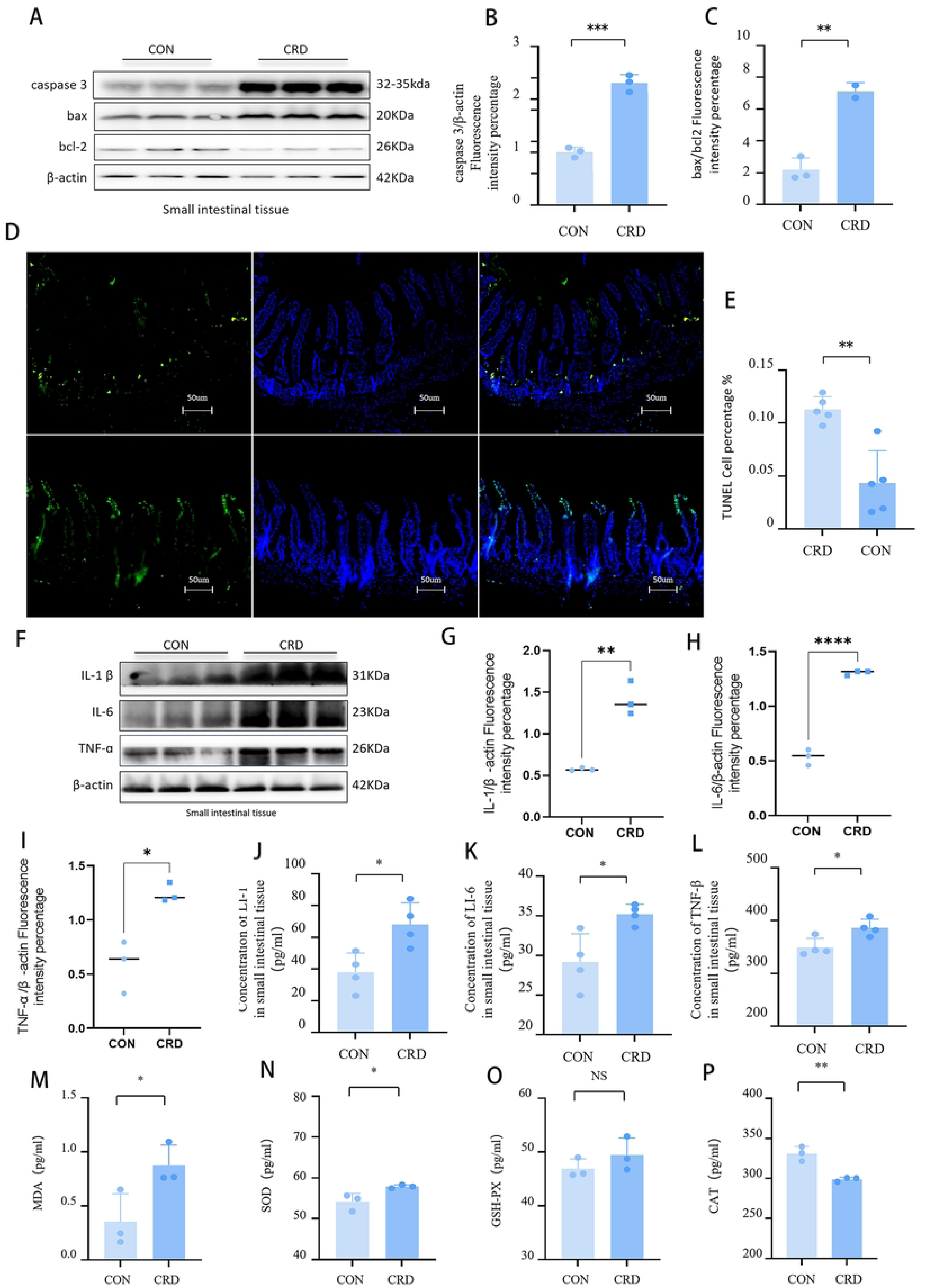
CRD promotes apoptosis, inflammation, and oxidative stress in intestinal tissue. (A) Representative western blot of apoptosis-related proteins: Cleaved Caspase-3, Bax, and Bcl-2. (B) Quantification of Cleaved Caspase-3 protein level. (C) Bax/Bcl-2 ratio. (D) Representative TUNEL staining images of intestinal tissue sections (scale bar = 50μm). (E) Quantification of TUNEL-positive cells. (F) Western blot of inflammatory cytokines: IL-1β, IL-6, and TNF-α. (G)Densitometric analysis of IL-1β protein levels normalized to β-actin, (H) IL-6, (I) TNF-α. (J) ELISA quantification of IL-1β in serum, (K) IL-6, (L)TNF-α. (M) MDA content in intestinal tissue. (N) CAT activity. (O) SOD activity. (P) GSH-PX activity. All data are presented as mean ± SEM(n=3). Statistical analyses were performed using unpaired Student’s t-test. CON, control group; CRD, circadian rhythm disruption group.

The overall trend of inflammatory factor Western blot was consistent with the ELISA results (Figure 6F-I), CRD group IL-1β levels were significantly increased (p=0.0086) (Figure 6J), IL-6 concentration in the CRD group (mean ± SEM: 15.2 ± 1.3 pg/mL) increased by 23.8% compared with the CON group (12.3 ± 1.1 pg/mL, p=0.0093() Figure 6K), CRD group TNF-α concentration increased by 9.3% compared with CON group (p=0.0030) (Figure 6L).

Malondialdehyde (MDA), as a lipid peroxidation marker, was significantly elevated in CRD group compared with CON group (p=0.0492)(Figure 6M). CAT activity in the CRD group (1.2 ± 0.2 U/mg protein) decreased by 90% compared with the CON group (12.0 ± 1.5 U/mg protein, p<0.05)(Figure 6N), Indicates impairment of the antioxidant enzyme defense system. In contrast, SOD activity was slightly elevated in CRD group (p=0.0373() Figure 6O).

In addition, the GSH-PX ratio was slightly higher in the CRD group(Figure 6P).

## 4. Discussion

Our findings demonstrate that circadian rhythm disruption significantly compromises intestinal barrier integrity, which is a key contributor to functional disorders such as Irritable Bowel Syndrome (IBS)^[14–16]^.

CRD experimental animals exhibited increased body weight with constant food intake ^[15]^, indicating reduced energy expenditure^[17]^. Anatomical observations revealed significant abdominal fat accumulation in CRD mice compared to CON mice, suggesting compensatory mechanisms like lipid metabolism abnormalities in C57 mice to counteract negative energy balance caused by circadian rhythm disruption^[18]^. Although this experiment did not address these mechanisms, we will investigate energy metabolism in subsequent studies. Additionally, treadmill analysis demonstrated highly dispersed daily exercise patterns and circadian rhythm delay effects. Behavioral rhythm abnormalities included reduced activity amplitude, random phase distribution, increased sleep fragmentation, and decreased dark cycle activity. These phenomena collectively reflect disrupted circadian rhythms^[19]^.

We validated the animal model at multiple levels. First, we examined the expression rhythms of core circadian clock genes (e.g., Clock, Bmal1, Per) at both central and peripheral levels. The imbalance between central and peripheral rhythms serves as a critical starting point for downstream pathological changes. CRD damages SCN output, characterized by phase shift and amplitude reduction in circadian gene rhythms^[20]^. Central system dysfunction leads to peripheral organ desynchronization, triggering systemic dysregulation that subsequently causes HPA axis dysfunction, elevated cortisol, inhibited melatonin synthesis, and disrupted circadian rhythms^[21]^. Core circadian clock gene analysis revealed that CRD intervention significantly disrupted molecular oscillation mechanisms in mouse hypothalamic suprachiasmatic nucleus (SCN), resulting in abnormal circadian transcriptional expression of key clock genes such as CLOCK, BMAL, CRY, and PER, manifested as amplitude reduction, phase delay, and rhythm disappearance. This indicates severe impairment of the core transcription-translation negative feedback loop (TTFL), particularly the abrupt decline in CLOCK gene expression at CT12, suggesting functional collapse that prevents effective activation of downstream clock-regulated genes^[22]^. In peripheral intestinal tissues, CRD also caused circadian gene dysregulation (e.g., CLOCK phase inversion, PER phase shift, and reduced BMAL1 amplitude). Despite compensatory attempts by peripheral oscillators, rhythm stability was lost—leading to central-peripheral clock decoupling, a key mechanism of CRD-induced gastrointestinal dysfunction^[11]^.

Secondly, in neuroendocrine studies, melatonin (MT) and 5-hydroxytryptamine (5-HT) exhibit opposite trends, phase delays and amplitude attenuation in secretion rhythms. These findings carry significant biological and clinical implications. As a circadian rhythm output signal, melatonin maintains mitochondrial homeostasis and regulates inflammatory responses. Abnormal signaling pathways constitute a critical component of circadian rhythm pathophysiology^[23]^. furthermore, melatonin possesses potentially antioxidant properties^[24]^, and the disorder of it secretion weakens cellular defense against oxidative damage and leads to intestinal epithelial barrier dysfunction^[25]^. Previous research indicates that melatonin abnormalities disrupt sleep-wake cycle stability, impair antioxidant capacity, and increase susceptibility to oxidative stress-related diseases^[26]^. 5-HT, serving as a precursor for melatonin synthesis, is highly associated with stress-sensitive neurotransmitter systems such as circadian rhythm systems^[27]^. From a neuroendocrine perspective, CRD causes delayed melatonin secretion phase and reduced amplitude^[28]^, This triggers a disorder of 5-HT secretion and a phase reversal^[29]^, reflecting severe impairment of neural regulation in core circadian timing systems. 5-HT disorders are also linked to mood fluctuations, appetite disturbances, gastrointestinal dysfunction, and exacerbate circadian rhythm disorders^[30]^. Neuroendocrine changes facilitate circadian rhythm disorder model establishment and enable future experiments. However, this study has limitations. First, we focused only on melatonin and serotonin secretion levels and rhythm changes, with mechanisms unclear, potentially involving altered gene expression, neurotransmitter interactions, and environmental factors. Future research should investigate these to understand disorder impacts. Second, as it used mouse models, caution is advised for human application; human studies should verify these phenomena.

Finally, behavioral abnormalities (such as autonomic activity rhythms and anxiety-like behaviors) are enhanced by circadian rhythm imbalance, increasing susceptibility to anxiety-like behaviors and depressive phenotypes^[31]^. In the open field experiment, the animal model showed a comprehensive impairment in behavioral aspects ^[32]^, including decreased total distance of activity; reduced exploration behavior and increased avoidance response to new environment^[33]^; significantly increased anxiety-like manifestations ^[34]^, extended stay time and reduced central exploration.

The intestinal barrier integrity, composed of tight junction proteins, mucus layer, and symbiotic microorganisms, serves to prevent pathogen translocation and enhance nutrient absorption^[35]^. The disruption of this integrity constitutes the direct pathological basis for IBS^[36]^. This study has demonstrated by various methods that CRD can impair the structural integrity, functional capacity and composition of symbiotic microbes of the intestinal barrier.

The fecal microbiota analysis (using 16S rRNA gene sequencing) revealed significant differences in gut microbiota composition between the CON group and CRD group. This finding slightly deviates from previous studies (where circadian rhythm disruption did not alter the microbiome of mice fed standard food)^[37]^, which we speculate may be related to the modeling approach. Our experimental design completely disrupted the circadian rhythms of mice, Future experiments will explore the differences and connections between different modeling methods.

The composition of the two groups was similar, with Bacteroidetes, Firmicutes, and Verrucomicrobia being dominant phyla. No statistically significant differences were observed between groups (P>0.05), indicating that CRD has not disrupted the core intestinal microbiota’s phylum-level homeostasis. The absence of significant differences in dominant phyla aligns with the resistance characteristics of the core intestinal microbiota: Bacteroidetes and Firmicutes serve as the foundation for homeostasis, compensating for environmental fluctuations in the CRD gut through metabolic redundancy to prevent collapse of the phylum-level structure^[38]^. This stability suggests that CRD’s interference with the microecosystem remains at a “compensatory” stage, providing potential opportunities for intervention.

LEfSe analysis revealed significant differences in microbial classification levels. The CON group was enriched with Bacteroidaceae (producing short-chain fatty acids like butyrate, which promote metabolic balance, reduce inflammation, and enhance intestinal tight junctions), Koserella (associated with metabolic balance), and Akkermansia muciniphila (maintaining gut barrier through mucosal repair and suppression of pro-inflammatory factors). These three groups collectively support intestinal homeostasis. In contrast, the CRD group exhibited enrichment with Prevotellaceae (associated with inflammation and metabolic disorders, capable of disrupting mucosal barriers and producing pro-inflammatory metabolites), δ-Proteobacteria (containing potential pathogenic bacteria like Campylobacter that may trigger systemic inflammation), and Alistipes (positively correlated with microbiota dysbiosis).Notably, Alistipes interferes with serotonin synthesis and exacerbates gut dysfunction. The overgrowth of these microbial groups may form a vicious cycle of “microbial dysbiosis-inflammatory exacerbation” in CRD patients.

The microbial diversity and structural separation were clearly defined. ANOSIM analysis revealed significant differences between groups, with 67.3% of OTUs overlapping between groups and 15.4% unique to the CRD group. However, Shannon index values showed substantial fluctuations, while the standard deviation of the CON group’s Shannon index was significantly lower than that of the CRD group, indicating greater microbial stability in the control group^[39]^. This phenomenon reflects “opportunistic colonization” under pathological conditions rather than the stability of beneficial diverse —— health-promoting microbiota (as evidenced by the low standard deviation in the CON group), which serves as the core indicator of microecological homeostasis. In contrast, the “diversity illusion” observed in the CRD group directly reflects microbial imbalance. β Principal Component Analysis (PCA), Principal Component Analysis of Ordination (PCoA), and Non-Metric Distance Scoring (NMDS) all demonstrated significant separation between the two groups (P<0.01), confirming that CRD drives the remodeling of microbial community structure.

At the macroscopic level, FITC-dextran in vivo fluorescence imaging technology visually reveals significant enhancement of intestinal permeability^[40]^. This abnormal permeability characteristic suggests that CRD may disrupt tight junction structures to facilitate macromolecular transmembrane transport, providing visual evidence for early pathological changes in “leaky gut syndrome”^[41]^. Notably, circadian rhythm-related pathologies (such as sleep deprivation and irregular shift work) are associated with increased intestinal barrier permeability^[42, 43]^. Furthermore, quantitative serum analysis of FITC-dextran further confirms this pathological presence^[44]^, highlighting the critical clinical significance of chronic circadian rhythm disorders (e.g., sleep-wake disturbances and improper shift schedules) that significantly elevate functional bowel disease risks, offering scientific support for clinical practice. Histopathologically, characteristic lesions including intestinal villous atrophy, aggravated inflammatory cell infiltration in the intestinal mucosal lamina propria, and shallow intestinal crypts are clearly observable^[45]^. CRD often triggers multiple synergistic factors, including intestinal villous structural abnormalities and chronic inflammation^[43]^. These structural changes interact with compromised intestinal barrier integrity to form a vicious cycle that collectively accelerates IBS progression. At the molecular level, key tight junction proteins (including ZO-1, Occludin, Claudin-1, and Claudin-3) exhibit widespread and significant downregulation in intestinal epithelial tissues. Taken together, these findings suggest that CRD severely impairs the physical barrier structure and function of the intestinal epithelial.

Extensive studies on relevant mechanisms have ultimately confirmed that CRD activates three key pathways: First, by releasing pro-inflammatory cytokines such as IL-1β, IL-6, and TNF-α, it exacerbates local tissue inflammation^[46]^. Additionally, the chronic inflammatory microenvironment induced by CRD significantly elevates reactive oxygen species (ROS) levels and weakens antioxidant enzyme activity^[47]^. Experimental results show increased malondialdehyde (MDA) levels (indicating enhanced lipid peroxidation) and inhibited catalase (CAT) activity (weakening antioxidant defense), demonstrating a pronounced oxidative stress state^[48, 49]^. Notably, SOD activity slightly increased in CRD groups, while the GSH-PX ratio was also marginally elevated, possibly reflecting compensatory regulatory mechanisms activated during early oxidative stress. Subsequent experiments will further explore these mechanisms through pathway inhibitors or overexpression studies, with focused efforts to investigate deeper signaling pathways to overcome current research limitations. Finally, activation of the intestinal epithelial cell apoptosis pathway——manifests as a significant increase in the Bax/Bcl-2 ratio, leading to enhanced mitochondrial membrane permeability, subsequent activation of Caspase-3 protease, and ultimately triggering apoptosis^[50]^. These three interconnected pathways form a positive feedback loop, collectively creating an “apoptosis-inflammation-oxidation” cascade amplification network that collaboratively damages intestinal barrier integrity, thereby initiating the development and progression of functional disorders.

This study has limitations requiring future refinement. The singular experimental model, using only male C57BL/6JNifdc mice without females or other strains like BALB/c, limits conclusion universality. Gender differences influence circadian rhythm regulation and gut microbiota; strains vary in light-induced CRD sensitivity. Findings cannot be generalized to females or other animal models, necessitating gender-matched and multi-strain validation.

Second, short CRD treatment duration fails to reflect chronic circadian disruption’s cumulative effects, as the experiment used only 21 days of light exposure without evaluating long-term impacts (e.g., 8-12 weeks) on intestinal barrier damage or reversibility after circadian restoration. Clinically, human circadian disorders like chronic shift work are chronic, so short-term results’ correlation with clinical scenarios requires verification. Future studies could design long-term CRD plus restoration experiments to clarify tissue damage’s temporal dependencies and reversibility.

Third, insufficient analysis of microbiota-barrier causality and functional aspects: The study showed CRD-induced microbial changes via 16S rRNA sequencing but omitted metagenomic analysis for functional insights. Fecal microbiota transplantation was not conducted to verify direct causality. Barrier evaluation focused solely on permeability and tight junction proteins, missing key indicators like mucus layer thickness and goblet cell count, potentially underestimating CRD’s impact on intestinal barriers.

The causal mechanism behind the apoptosis-inflammation-oxidative stress cascade is still not fully understood. CRD concurrently triggers apoptotic, inflammatory, and oxidative stress pathways; however, no intervention experiments were performed to confirm the upstream-downstream relationships, such as whether oxidative stress occurs before inflammation or whether microbiota imbalance worsens inflammation through apoptosis. This absence of causal validation makes the “CRD impairs intestinal barrier via dysbiosis and cascade reaction” mechanism incomplete, necessitating further confirmation through pathway blocking experiments. Additional experiments will be carried out to further validate the pathway blocking test.

## 5. Conclusion

This study established a circadian rhythm disruption (CRD) model in male C57BL/6 mice via 21-day continuous light exposure, and confirmed that CRD impairs intestinal barrier integrity through two key pathways:1.gut microbiota dysbiosis (e.g., decreased Prevotellaceae and Alistipes, increased Bacteroidaceae and Akkermansia); 2.activation of the apoptosis-inflammation-oxidative stress cascade (e.g., elevated Caspase-3 activity, increased pro-inflammatory cytokines IL-1β/IL-6, and enhanced lipid peroxidation marker MDA). The observed intestinal barrier damage (increased permeability, downregulated tight junction proteins, shortened villi) provides a mechanistic link between CRD and irritable bowel syndrome (IBS). These findings identify gut microbiota and the apoptosis-inflammation-oxidative stress cascade as potential therapeutic targets for IBS. Future studies should validate the findings in female mice and other strains, explore long-term CRD effects, and clarify the causal relationship between microbiota and barrier damage via fecal microbiota transplantation.”

## 6. Statement

### 1. Ethics Statement

All procedures were approved by the Ethics Committee of PLA Rocket Force Characteristic Medical Center (Approval No.: KY2024040) and conducted in accordance with the Guide for the Care and Use of Laboratory Animals (8th ed., National Research Council, USA), the NIH Guidelines for the Care and Use of Laboratory Animals, and the ARRIVE Guidelines. Euthanasia via cervical dislocation was performed under anesthesia to minimize animal suffering, adhering to humane endpoints.

## Acknowledgments

We thank the technical support from the Laboratory of the PLA Rocket Force Characteristic Medical Center for assistance with animal monitoring and sample processing.Chang-Zheng Li takes responsibility for the integrity of the work as a whole, from inception to published article. Tai-Wei Zhang performed the research, Tai-Wei Zhang and Jia-Chun Song collected the data, Tai-Wei Zhang, Ningbo Hao and Jia-Chun Song analysed the data, all authors have contributed to the study design and manuscript revision. All authors read and approved the final manuscript and consented to publish this manuscript.

## Data and material availability

The raw data that support the findings of this study are available from the corresponding author (Chang-Zheng Li) upon reasonable request. Raw Western blot images are provided in Supplementary Materials.

## Author Contributions

Tai-Wei Zhang (TWZ): Conceptualization, Methodology, Investigation, Data Curation, Writing - Original Draft. Jia-Chun Song (JCS): Investigation, Data Curation, Writing - Review & Editing. Ning-Bo Hao (NBH): Data Curation, Writing - Review & Editing. Ming-Yue Qu (MYQ): Conceptualization, Writing - Review & Editing. Bao-Shi Guo (BSG): Conceptualization, Writing - Review & Editing. Chang-Zheng Li (CZL): Conceptualization, Supervision, Funding Acquisition, Writing - Review & Editing, Corresponding Author. All authors read and approved the final manuscript.

## Declaration of Generative AI and AI-Assisted Technologies

No generative artificial intelligence (AI) tools (e.g., ChatGPT, Jasper) were used in the writing, editing, or data analysis of this manuscript. All content reflects the authors’ original work and critical thinking.

---

**Table 1.** Standard curve preparation concentration gradient.

| Concentration gradient (ng/ml) | Preparation method |
| --- | --- |
| 5000 | 1ul 5mg/mlFITC-dextran+999ul 1*pbs |
| 500 | 100ul 5000ng/mlFITC-dextran+900ul 1*pbs |
| 50 | 100ul 500ng/mlFITC-dextran+900ul 1*pbs |
| 5 | 100ul 50ng/mlFITC-dextran+900ul 1*pbs |
| 0.5 | 100ul 5ng/mlFITC-dextran+900ul 1*pbs |
| 0.05 | 100ul 0.5ng/mlFITC-dextran+900ul 1*pbs |
| 0 | 1*pbs |

**Table 2.** Antibody List.

| Antibody Name | Manufacturer | Dilution | Catalog Number |
| --- | --- | --- | --- |
| BAX Polyclonal antibody | proteintech | 1:1500 | 50599-2-Ig |
| Bcl2 Polyclonal Antibody | proteintech | 1:1000 | 26593-1-AP |
| Beta Actin Polyclonal Antibody | proteintech | 1:1500 | 20536-1-AP |
| Anti-PER1 antibody | Proteintech | 1:1000 | 13463-1-AP |
| TNF Alpha Rabbit Polyclonal antibody | Proteintech | 1:500-1:2000 | 26405-1-AP |
| IL-1beta Rabbit Polyclonal antibody | Proteintech | 1:500-1:3000 | 26048-1-AP |
| GAPDH Monoclonal antibody | Proteintech | 1:5000 | 60004-1-Ig |
| Anti-Caspase-3 antibody [EPR18297] | Abcam | 1:2000 | ab184787 |
| Anti-ZO1 tight junction | Abcam | 1:1000 | ab96587 |
| Anti-Claudin 3 antibody [EPR28731-2] | Abcam | 1:800 | ab317319 |
| Anti-Occludin antibody [EPR20992] | Abcam | 1:1000 | ab216327 |
| Anti-Claudin 1 antibody [EPR25359-48] | Abcam | 1:1000 | ab307692 |
| Anti-KAT13D/CLOCK antibody | Abcam | 1:1000 | ab3517 |
| Anti-BMAL1 antibody | Abcam | 1:1000 | ab230822 |
| Anti-Cryptochrome 1/CRY1 antibody | Abcam | 1:1000 | ab104736 |
| Anti-IL-6 antibody [EPR23819-103] | Abcam | 1:1000 | ab290735 |

## Abbreviation list

CRD: Circadian Rhythm Disruption
IBS: Irritable Bowel Syndrome
FITC: Fluorescein Isothiocyanate
WB: Western Blotting
TUNEL: TdT-mediated dUTP Nick-End Labeling
ELISA: Enzyme-Linked Immunosorbent Assay
MT: Melatonin
5-HT: 5-Hydroxytryptamine
SOD: Superoxide Dismutase
GSH-PX: Glutathione Peroxidase
MDA: Malondialdehyde
CAT: Catalase
OTU: Operational Taxon Units
PCA: Principal Component Analysis
PCoA: Principal Coordinate Analysis
NMDS: Non-Metric Multidimensional Scaling
SPF: Specific Pathogen-Free
ZT: Zeitgeber Time
CT: Circadian Time

